# Comparing the folding landscapes of evolutionarily divergent procaspase-3

**DOI:** 10.1101/2022.01.12.476064

**Authors:** Liqi Yao, A. Clay Clark

## Abstract

All caspases evolved from a common ancestor and subsequently developed into two general classes, inflammatory or apoptotic caspases. The caspase-hemoglobinase fold has been conserved throughout nearly one billion years of evolution and is utilized for both the monomeric and dimeric subfamilies of apoptotic caspases, called initiator and effector caspases, respectively. We compared the folding and assembly of procaspase-3b from zebrafish to that of human effector procaspases in order to examine the conservation of the folding landscape. Urea-induced equilibrium folding/unfolding of procaspase-3b showed a minimum three-state folding pathway, where the native dimer isomerizes to a partially folded dimeric intermediate, which then unfolds. A partially folded monomeric intermediate observed in the folding landscape of human procaspase-3 is not well-populated in zebrafish procaspase-3b. By comparing effector caspases from different species, we show that the effector procaspase dimer undergoes a pH-dependent conformational change, and that the conformational species in the folding landscape exhibit similar free energies. Together, the data show that the landscape for the caspase-hemoglobinase fold is conserved, yet it provides flexibility for species-specific stabilization or destabilization of folding intermediates resulting in changes in stability. The common pH-dependent conformational change in the native dimer, which yields an enzymatically inactive species, may provide an additional, albeit reversible, mechanism for controlling caspase activity in the cell.

## Introduction

The caspase family is an attractive model to examine the evolution of protein folding landscapes. Caspase genes predate multicellularity and are widely present in eukaryotes (1). Caspases are thought to have evolved from a common ancestral protein and later divided into two general classes, inflammatory or apoptotic caspases, through gene and genome duplication events (2). The apoptotic caspases further evolved into two subfamilies, the initiator or effector caspases, and the ancestral effector caspase later evolved into three modern proteins: caspases-3, -6, and -7. Importantly, the caspase-hemoglobinase fold has been conserved throughout the nearly one billion years of evolution leading to the modern enzyme subfamilies.

Caspases exist as latent zymogens in cells and are activated upon induction of apoptosis or the inflammatory response (3). The zymogens of initiator caspases exist as protomers, and they are activated by dimerization, while the zymogens of effector caspases exist as stable, yet inactive dimers, and they are activated by cleavage of the intersubunit linker (4). Each mature caspase is a homo-hetero-dimer ((LS)_2_), containing two protomers (Figure 1A). The large (L) and small (S) subunits comprise a protomer and fold as the single domain consisting of a six-stranded β-sheet core surrounded by five *α*-helices, with a single active site (Figure 1B). In effector caspases, cleavage of the intersubunit linker that connects the large and small subunits results in rearrangement of several loops in the active site, allowing the substrate-binding pocket to form (2). It is not yet clear how the subfamilies of dimers *versus* monomers evolved from the same conserved protein fold, called the caspase-hemoglobinase fold (2), but dimerization is a key regulatory mechanism controlling apoptosis (3,4).

**Figure 1.**
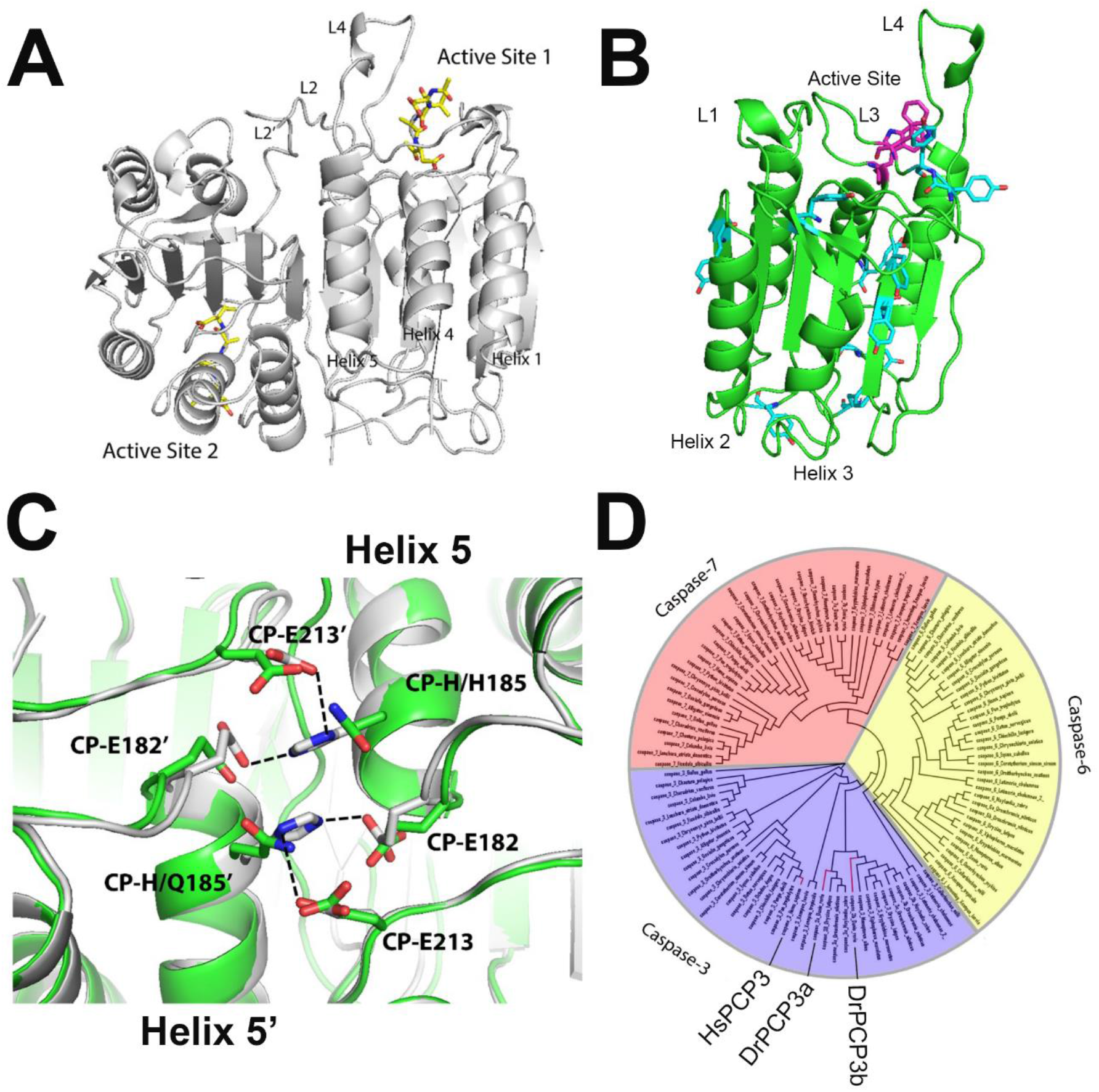
Caspase structure and phylogeny. A. Structure of human caspase-3 dimer is shown in grey (PDB ID: 1CP3). Inhibitors are shown in yellow, and active site loops L2, L2’ and L4 are labeled. B. Caspase protomer with active site tryptophans (magenta) and tyrosines (cyan) labeled. Active site loops L1, L3, and L4 are shown, and helices 2 and 3 are labeled. C. Comparison of contacts across the dimer interface (helix 5) for zebrafish caspase-3b (green) (modeled from human caspase-3) and human caspase-3 (grey) (PDB ID: 1CP3). CP refers to the common position numbering system as described in the text. Dashed lines show hydrogen bonds. D. Phylogenetic tree of effector caspases. Caspase-3, -6 and -7 are marked in purple, yellow and red, respectively. Human caspase-3 (HsPCP3), zebrafish caspase-3a (DrPCP3a) and -3b (DrPCP3b) are labeled and shown by the red lines.

We showed previously that human procaspase-3 unfolds via a four-state equilibrium process in which the native dimer (N_2_) isomerizes to a dimeric intermediate (I_2_), and the dimeric intermediate dissociates to a partially folded protomer (I) which then unfolds (5). We also showed that the dimer of human procaspase-3 undergoes a pH-dependent dissociation below pH 5, such that the protein is monomeric at pH 4 (6). Dissociation of the native ensemble at lower pH is most likely due to disruption of electrostatic interactions across the dimer interface between CP-E182, CP-H185, and CP-E213 in helix 5, and the corresponding residues from the second protomer (Figure 1C). We note that “ CP” refers to the “ common position” naming system, described previously, that is utilized to denote common sequence positions of all caspases (7). In addition, we recently described the common position nomenclature for the proteins examined here (8) and show the sequence alignment in Supplemental Figure S1.

While human procaspase-3 has been extensively studied for its equilibrium and kinetic folding properties (5,6,9,10), we recently examined the evolution of the folding landscape of effector caspases (11) and showed that the landscape was established with the common ancestor of effector caspases more than 650 million years ago. Evolutionary changes in the relative populations of the dimeric (I_2_) versus monomeric folding intermediates (I) in human procaspases-3, -6, -7 appear to provide tremendous flexibility in species-specific evolution of the caspase dimer. From the previous studies, however, only procaspase-3 undergoes a pH-dependent dissociation, but all caspase dimers from human effector caspases (caspase-3, -6, and -7) undergo a reversible pH-dependent conformational change from the native dimer (N_2_) to the enzymatically inactive dimeric intermediate (I_2_) (11), which may provide additional control of enzyme activity in the cell. It was not clear, however, whether the data reflect the properties of all effector caspases or is a more specific for human caspases. That is, the conservation of the folding landscape and potential species-specific evolutionary changes are not known due to a dearth of studies that characterize the folding of effector caspases from humans or other species.

To further understand the evolution of effector caspases, we examined the folding and assembly of procaspase-3 from zebrafish (*Danio rerio*). Zebrafish caspases are an excellent model to further our understanding of the caspase folding landscape because zebrafish experienced a whole genome duplication around 300 million years ago (12), resulting in two copies of the caspase-3 gene, called caspase-3a and caspase-3b. Based on a phylogenetic analysis (Figure 1D and Supplemental Table S1), human caspase-3 and zebrafish caspases-3a and -3b are evolutionarily close, compared to the caspase-6 and -7 subfamilies. Human caspase-3 has 56% amino acid sequence identity compared to the two zebrafish caspase-3 proteins, while the two zebrafish caspase-3 proteins have a 59% sequence identity compared to each other (13).

We examined the urea-induced equilibrium folding and assembly of the zymogen of zebrafish caspase-3b over a broad pH range, from pH 5 to 8.5. We show that the folding landscape is similar to that described previously for human procaspase-3 (6) and procaspase-6 (11) in that the native dimer isomerizes to a dimeric intermediate, which then unfolds. The partially folded monomeric intermediate observed in human procaspase-3 is not well populated in the ensemble of zebrafish procaspase-3b, similarly to human procaspase-6. In contrast, however, one observes an additional dimeric intermediate at neutral pHs in the ensemble of zebrafish procaspase-3b. Finally, unlike human procaspase-3, zebrafish procaspase-3b is a dimer at all pHs examined. Collectively, the data suggest that evolutionary changes in a conserved folding landscape result in species-specific selection of folding intermediates available within the landscape.

## Results and Discussion

For all experiments described here, we used a catalytically inactive mutant of zebrafish procaspase-3b, CP-C117S since the wild-type procaspase can undergo self-cleavage during heterologous expression (8,14). Unfortunately, we were unable to purify zebrafish procaspase-3a in sufficient quantities for the biophysical studies described here, so we have focused on comparing zebrafish procaspase-3b with human procaspase-3, henceforth called DrPCP3b (*Danio rerio* procaspase-3b) and HsPCP3 (*Homo sapiens* procaspase-3). Generally, the DrPCP3b protomer has 293 amino acids and a molecular weight of 32,983 Da, including the LEH_6_ C-terminal sequence used for purification. In addition, DrPCP3b contains two tryptophan residues located in the active site, in the same positions as those in HsPCP3 (Supplemental Figure S1). Finally, DrPCP3b contains ten tyrosine residues that are well-distributed throughout the structure (Figure 1B and Supplemental Figure S1). As described previously (15), we examined changes in tertiary structure in the presence of urea by monitoring changes in fluorescence emission, after excitation at 280 or 295 nm. Excitation at 280 nm follows the tyrosinyl and tryptophanyl fluorescence emission, while excitation at 295 nm follows the emission of tryptophanyl residues. We also monitored changes in secondary structure by circular dichroism (CD).

When DrPCP3b is excited at 280 nm, the protein exhibits a peak in fluorescence emission at 338 nm (Figure 2A), which is similar to that of HsPCP3 (emission maximum of 335 nm (16)). When unfolded in 8 M urea-containing buffer, however, the fluorescence emission is red-shifted to ∼347 nm (Figure 2A). When the tryptophan residues were examined using an excitation wavelength of 295 nm, the data show that the two tryptophans in the active site are in a hydrophilic environment, with fluorescence emission maximum of 342 nm (Figure 2B), again similar to HsPCP3 (emission maximum of 340 nm (16)). The fluorescence emission of the tryptophan residues is also red-shifted to 352 nm when the protein is unfolded in 8 M urea-containing buffer (Figure 2B). Similarly, DrPCP3b exhibits well-formed secondary structure, as measured by far-UV CD, which is disrupted in 8 M urea-containing buffer (Figure 2C).

**Figure 2.**
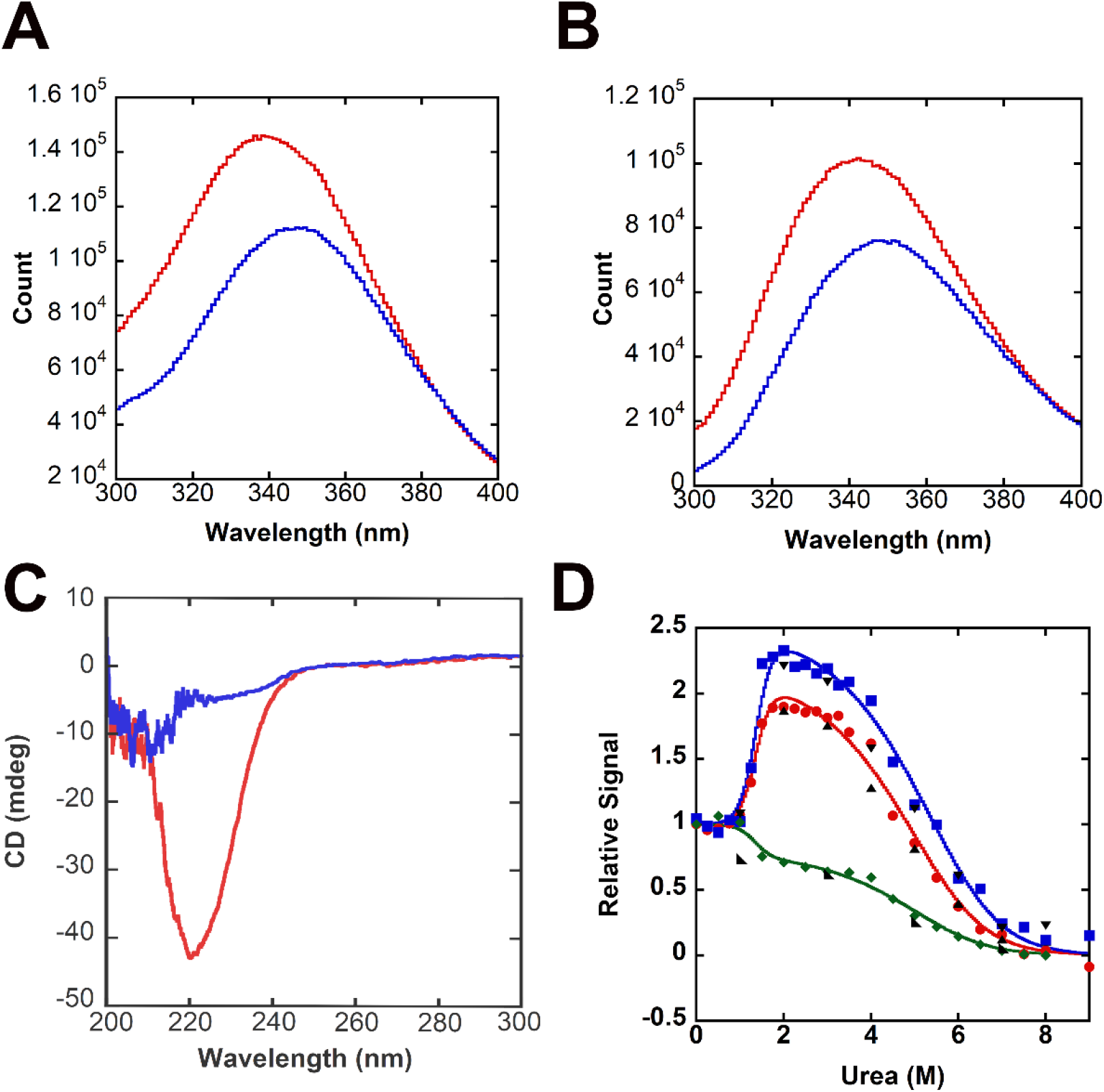
Spectroscopic properties of zebrafish procaspase-3b (CP-C117S). A. Fluorescence emission scan of DrPCP3b following excitation at 280 nm in buffer containing zero urea (red) or 8 M urea (blue). B. Fluorescence emission scan of DrPCP3b following excitation at 295 nm in buffer containing zero urea (red) or 8 M urea (blue). C. Circular dichroism (CD) far-UV scan of DrPCP3b in buffer containing zero urea (red) or 8 M urea (blue). D. Representative equilibrium unfolding/folding of DrPCP3b at pH 7 by fluorescence emission (average emission wavelength) following excitation at 280 nm 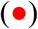, 295 nm 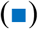, and CD at 224 nm 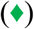. Refolding data for fluorescence emission (280 nm (▲) and 295 nm (▼)) and CD (◣) show that folding is reversible. Solid lines represent global fits to a three-state folding model, as described in the text.

### Equilibrium unfolding of DrPCP3b

We examined the equilibrium unfolding of DrPCP3b as a function of urea concentration, from 0 to 9 M, and between pH 5 and pH 8.5, as described in Materials and Methods. Because DrPCP3b is a homodimer, we also examined the effect of protein concentration on equilibrium unfolding, using a protein concentration range of 0.5-4 μM for fluorescence emission experiments or 2 μM to 8 μM for CD. Representative data for the unfolding of DrPCP3b at pH 7 and 4 μM protein are shown in Figure 2D and are described in more detail below.

For both the fluorescence emission and CD data, one observes a pre-transition between 0 M and ∼1.5 M urea, which shows little to no change in the signal of the native dimer. Following the pre-transition, the signal changes cooperatively between ∼1.5 M and ∼3 M urea. While the fluorescence emission increases in this transition, the CD signal decreases. In addition, the relative fluorescence emission from excitation at 295 nm is higher in ∼3 M urea than that of protein excited at 280 nm, and they are both higher than the CD signal. At higher urea concentrations, one observes a second transition where both fluorescence emission and CD signals decrease cooperatively. The protein is unfolded at urea concentrations >7 M. Collectively, the data show that a partially unfolded intermediate conformation is well-populated between ∼3 and 5 M urea and that the fluorescence emission of the intermediate is less quenched compared to that of the native conformation. Variations in the protein concentration show that there is little to no protein concentration dependence to the first transition, from 0 M to ∼3 M urea (Supplemental Figures S2 and S3). In contrast, the mid-point of the second transition increased to higher urea concentrations as the protein concentration was also increased. Overall, the data at pH 7 (Figure 2D) show that DrPCP3b undergoes a three-state equilibrium unfolding process in which the native dimer (N_2_) isomerizes to a dimeric intermediate (I_2_), which then unfolds to yield the unfolded state (U).

Together, the experimental procedure yields at least eleven data sets, and we fit the data globally to a three-state equilibrium unfolding model in which the native dimer isomerizes to a dimeric intermediate which then unfolds (see equation 2). The results of the fits are shown as the solid lines in the figures. At pH 7, the global fits show that the first isomerization (N_2_⇄I_2_) occurs with a free energy of ΔG_1_°_conf_ =4.4±0.1 kcal mol^-1^ and that the dimeric intermediate unfolds (I_2_⇄2U) with ΔG_3_°_conf_ =11.5±2.1 kcal mol^-1^, with a total conformational free energy of 15.9 kcal mol^-1^ at pH 7 and 25 °C (Table 1). The associated m-values (m_1_=3.20±0.20 kcal mol^-1^ M^-1^; m_3_=0.87±0.10 kcal mol^-1^ M^-1^) show that the formation of the dimeric intermediate (I_2_) is correlated with substantial exposure of buried surface area relative to the native dimer (N_2_). Together, the data are similar to those previously determined for HsPCP3 (6), although all conformations of DrPCP3b exhibit lower conformational free energies compared to HsPCP3. For example, HsPCP3 has a total conformational free energy of 25.8 kcal mol^-1^ at pH 7 and 25 °C, with a free energy change of 8.3±1.3 kcal mol^-1^ for the formation of the dimeric intermediate, I_2_. In comparison to HsPCP6 and HsPCP7, DrPCP3b is more similar to HsPCP7, which has a total conformational free energy of 15.4 kcal mol^-1^ (11). In contrast with HsPCP3, however, the monomeric intermediate is not well-populated in DrPCP3b, so the unfolding data are more similar to that of HsPCP6, where the second unfolding transition reflects the unfolding of the dimer to the unfolded conformation (11). The lower conformational free energy of DrPCP3b compared to that of HsPCP3 (∼5 kcal mol^-1^) is most likely due to the lower stability, and thus lower fractional population, of the monomeric folding intermediate.

**Table 1.** Summary of free energy and m-values for unfolding DrPCP3b at all pHs.

| pH | $\Delta G_1$<br>(kcal<br>mol<sup>-1</sup>) | $m_1$ (kcal<br>mol<sup>-1</sup>M<sup>-1</sup>) | $\Delta G_2$<br>(kcal<br>mol<sup>-1</sup>) | $m_2$ (kcal<br>mol<sup>-1</sup>M<sup>-1</sup>) | $\Delta G_3$<br>(kcal<br>mol<sup>-1</sup>) | $m_3$ (kcal<br>mol<sup>-1</sup>M<sup>-1</sup>) | $\Delta G_{\text{total}}$<br>(kcal<br>mol<sup>-1</sup>) | $m_{\text{total}}$<br>(kcal<br>mol<sup>-1</sup>M<sup>-1</sup>) |
| --- | --- | --- | --- | --- | --- | --- | --- | --- |
| 5 |  |  | 0.5±0.2 | 1.4±0.2 | 14.4±1.7 | 1.4±0.1 | 14.9±1.9 | 2.8±0.2 |
| 5.5 |  |  | 2.3±0.5 | 2.6±0.52 | 13.1±1.5 | 1.1±0.1 | 15.4±2 | 3.7±0.6 |
| 6 | 3.8±2 | 2.8±0.5 | 0.9±0.3 | 2.3±0.4 | 12.5±0.4 | 0.92±0.1 | 17.2±2.7 | 6.1±1.0 |
| 6.5 | 5.3±0.8 | 2.9±0.4 | 0.3±0.1 | 1.8±0.5 | 11.8±2.4 | 1.37±0.1 | 17.4±3.3 | 6.1±1 |
| 7 | 4.4±0.1 | 3.2±0.2 |  |  | 11.5±2.1 | 0.9±0.1 | 15.9±2.2 | 4.1±0.3 |
| 7.5 | 5.4±0.5 | 3.4±0.3 |  |  | 12.6±0.7 | 1.0±0.1 | 18.5±1.2 | 4.4±0.4 |
| 8 | 4.5±0.5 | 2.8±0.3 |  |  | 12.5±0.3 | 0.9±0.1 | 17.5±0.8 | 3.7±0.4 |
| 8.5 | 2.6±0.2 | 2.6±0.2 |  |  | 11.3±0.4 | 0.7±0.1 | 13.9±0.6 | 3.3±0.3 |

### pH effects on the native dimer

We showed previously that the dimer of HsPCP3 (6) and of HsPCP6 (11) undergoes a pH-dependent conformational change to an inactive dimeric conformation (I_2_), with pKa∼5.7. In the case of HsPCP3, the dimer then dissociates below pH∼5 such that the protomer is fully populated at pH 4 in the absence of urea (6). We performed similar experiments for DrPCP3b to determine whether the pH effects are a common feature of the procaspase-3 dimer. In this case, we examined the urea-induced equilibrium unfolding of DrPCP3b between pH 5 and pH 8.5, and the results are shown in Supplemental Figures S2 and S3. In each panel, we also show results of refolding protein from 9 M urea-containing buffer, which demonstrate that folding is reversible over the entire pH range. We note that we were unable to conduct experiments below pH 5 because, unlike HsPCP3 and HsPCP6, DrPCP3b precipitates at low pH.

Initially, while performing urea-induced equilibrium unfolding experiments at several pHs, we observed that the fluorescence emission of the native dimer depends on the pH of the buffer. As shown in Figure 3A, at pH >6, the average emission wavelength (AEW) of DrPCP3b is 347 nm or 343 nm, respectively, when excited at 295 nm or 280 nm. One observes that the AEW is blue-shifted to lower values at pH<6, indicating that the aromatic amino acids are in a more hydrophobic environment. We examined the protein by size exclusion chromatography over the same pH range, and the data show a single peak at ∼70 kDa (Figure 3B), demonstrating that the protein remains dimeric at pH 5 even though the AEW is blue-shifted. Based on fits of the data in Figure 3A, as described previously (6,11), we estimate the pKa of the transition to be ∼5.4. Collectively, the data show that the dimer of DrPCP3b undergoes a pH-dependent transition with a pKa similar to that observed of HsPCP3 and of HsPCP6, and the transition correlates with a change in environment for the aromatic residues. Because the two tryptophans are in the active site, the results suggest that one or more active site loops comprise the pH-dependent transition.

**Figure 3.**
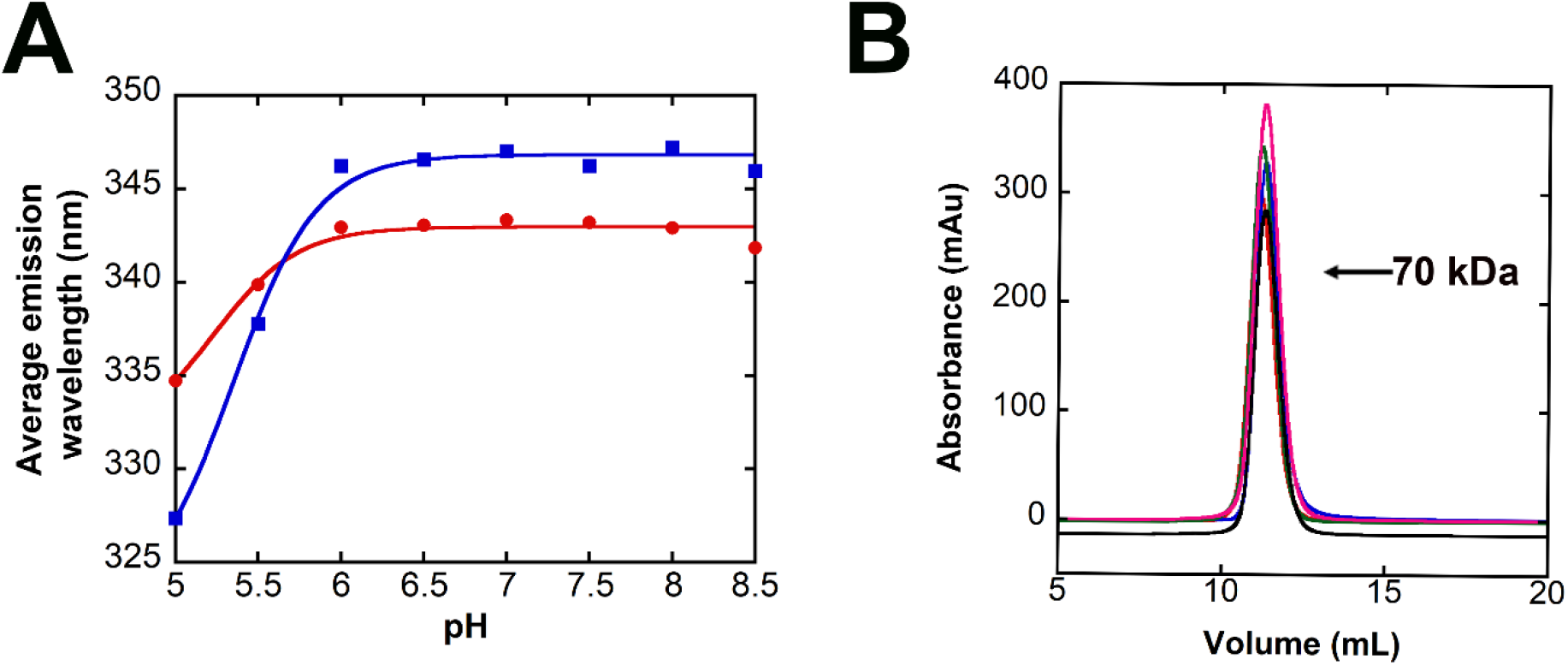
Effect of pH on the DrPCP3b dimer. A. Average emission wavelength following excitation at 280 nm (red) or 295 nm (blue) versus pH for DrPCP3b. Solid lines represent fits of the data to determine the pKa of the transition, as described in the text. B. Size exclusion chromatography of DrPCP3b over the pH range shown in panel A. Protein was incubated citrate buffer at pH 5 (red), pH 6 (blue), and pH 6.5 (green), and in phosphate buffer at pH 7.5 (black) and pH 8 (magenta). The column was standardized as described in the text to determine the molecular weight of 70 kDa for the single peak in the chromatograms.

### pH effects on equilibrium unfolding

As described above for experiments at pH 7 (Figure 2D), the results for urea-indued equilibrium unfolding of DrPCP3b over the pH range of 5-8.5 are summarized in Figure 4, and all data are shown in Supplemental Figures S2 and S3. Between pH 7 and 8.5, the urea-induced equilibrium unfolding data are well described by the three-state folding mechanism (N_2_⇄I_2_⇄2U, see equation 2). The protein exhibits a pre-transition between 0 M and ∼1.5 M urea, which is followed by a cooperative transition to a state with higher relative fluorescence emission and less secondary structure (Figure 2D and Supplemental Figure S3). At higher urea concentrations (>∼4 M), one observes a second cooperative transition that results in the unfolded protein. The folding intermediate that is populated in the first transition (∼3 M urea) is shown to be dimeric since the mid-point of the second transition is dependent on the protein concentration.

**Figure 4.**
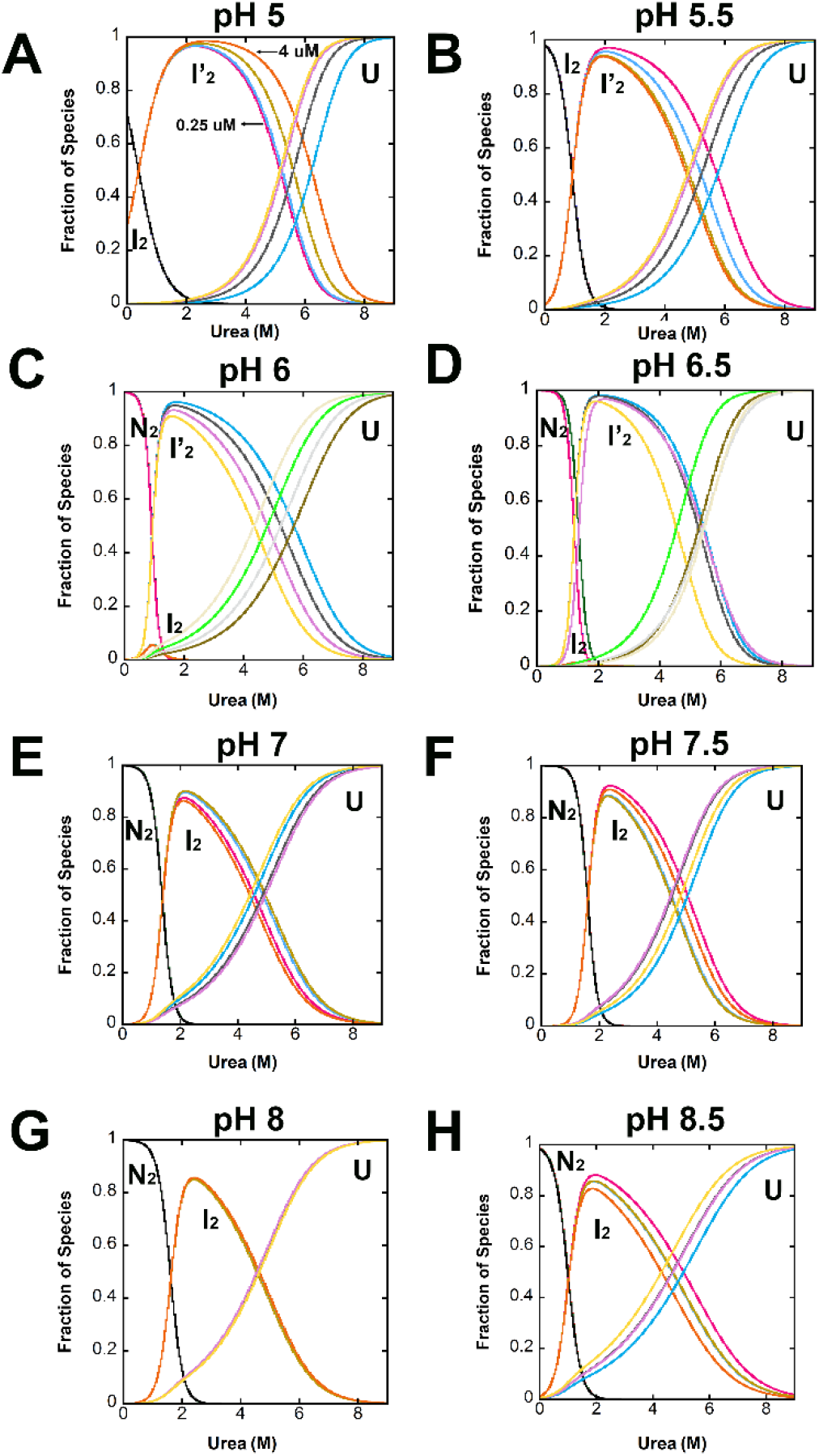
Fraction of species as a function of urea concentration over the pH range of 5 to 8.5. The fractions of species versus urea concentration at pH 5 (A), pH 5.5 (B), pH 6 (C), pH 6.5 (D), pH 7 (E), pH 7.5 (F), pH 8 (G) and pH 8.5 (H) were calculated from global fits of the equilibrium folding/unfolding data shown in Supplemental Figures S2 and S3, the parameters shown in Table 1, and the protein concentrations of 0.5 µM, 1 µM, 2 µM, and 4 µM. Each species, N_2_, I_2_, I_2_’, U, is labeled in the figures. N_2_ refers to the native dimer of DrPCP3b, while I_2_’ and I_2_ refer to partially folded intermediate conformations of the dimer, and U refers to the unfolded state of the protein.

For each pH, we calculated the equilibrium distribution of species over the urea concentration range of 0-9 M by using the values of the free energies, the cooperativity indices determined for each transition, and four protein concentrations (0.5, 1, 2, and 4 µM). Representative data for pH 5-8.5, corresponding to the data from Supplemental Figures S2 and S3, are shown in Figure 4, and the conformational free energies and m-values are described more fully below (Figure 5). As the protein is unfolded in urea-containing buffers below pH 7 (Supplemental Figure S2), one observes that the midpoint of the first transition shifts to lower urea concentrations such that the transition is no longer observed at pH 5. The shift in the mid-point of the first unfolding transition suggests that the native dimer is less stable at pH<7 (Supplemental Figures S2 and S3). Secondly, at intermediate pH values (pH 6.5 and pH 6), one observes an additional transition in the data. In this case, the dimeric intermediate (I_2_) is well-populated in ∼1.5 M urea, then the dimer undergoes an additional conformational change prior to unfolding (Figure 4 and Supplemental Figure S2). The transition is characterized by a decrease in fluorescence emission, compared to the dimeric intermediate (I_2_), but without a further loss in secondary structure. The protein concentration dependence to unfolding shows that the dimer dissociates and unfolds above ∼4 M urea, similar to the data at higher pHs. The results show that the transition between the dimeric intermediate, I_2_, and unfolded protomer is not affected by the change in pH. In this case, the mid-point for the transition is ∼5-6 M urea, depending on the protein concentration. Overall, we interpret these results to show that the population of native dimer (N_2_) decreases below pH 6 and is replaced by the dimeric intermediate (I_2_) as the predominant species in the absence of urea. The dimeric intermediate, I_2_, is populated maximally at pH 5.5 and decreases at pH 5 such that the native ensemble is a mixture of the two dimeric intermediates, I_2_ and I_2_’. Presumably, the dimeric intermediate I_2_’ would become the predominant species at pH below 5, based on the trend observed in the data. The second dimeric intermediate (I_2_’) is characterized by a somewhat lower fluorescence emission compared to the I_2_ conformation but a higher relative fluorescence compared to the native dimer (N_2_). The secondary structure appears to be similar between the two dimeric intermediates, I_2_ and I_2_’, since we observe no change in the CD signal from ∼1.5 M to ∼4 M urea that would distinguish the two conformations.

**Figure 5.**
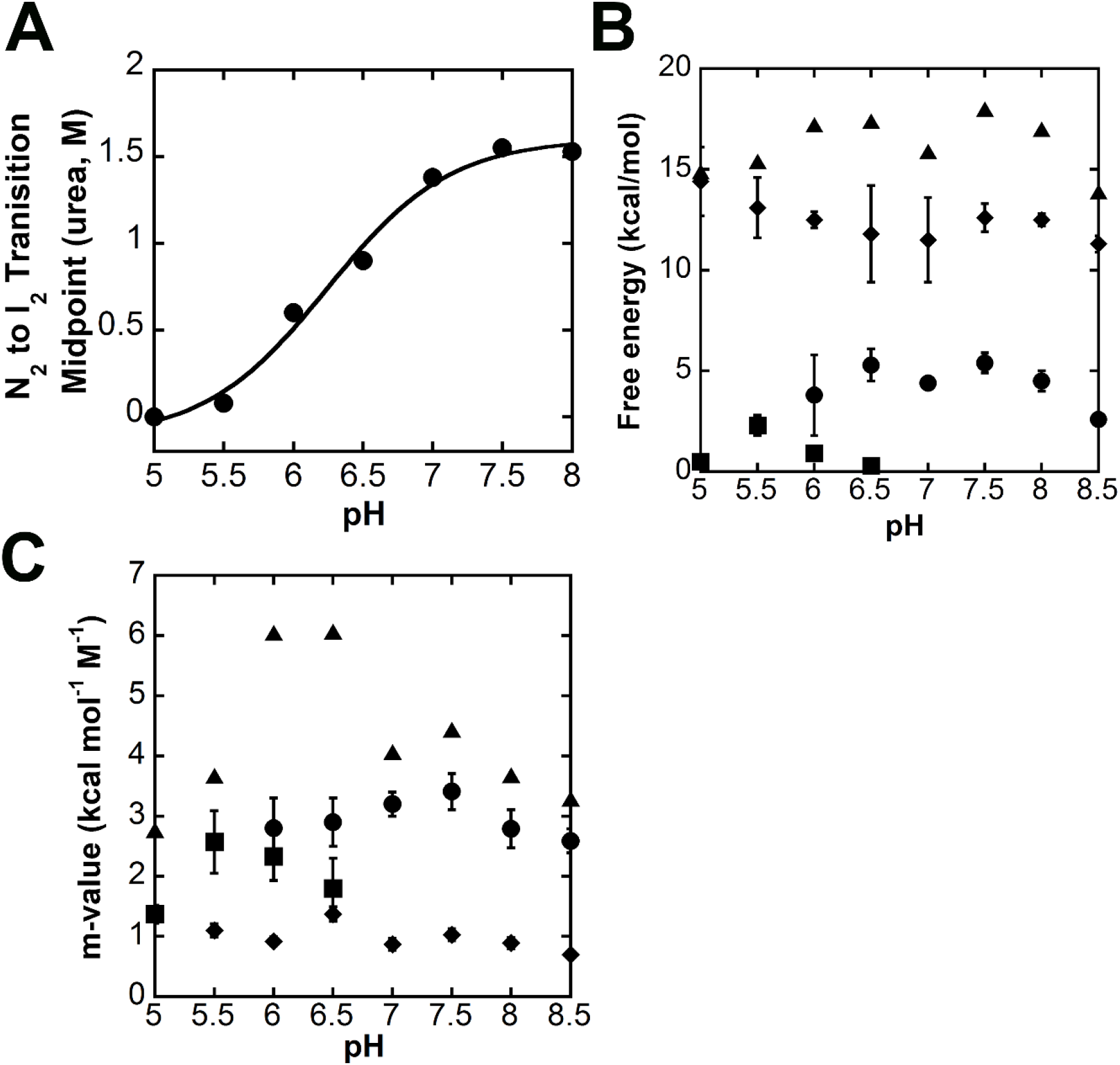
Conformational free energy of DrPCP3b versus pH. A. The midpoint of the transition between native dimer (N_2_) and the first dimeric folding intermediate (I_2_) (●). The solid line represents a fit of the data to determine pKa, as described in the text. B. Free energies for 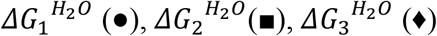 and 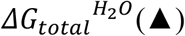 *versus* pH. C. Cooperativity indices, m_1_ ((●), m_2_ (■), m_3_ (♦) and m_total_ (▲) *versus* pH. Data for panels B and C were determined from global fits of the data shown in Supplemental Figure S2 and S3 to the models shown in equations 2 and 3, as described in the text. The error bars show the standard error obtained from the global fits.

We suggest that both dimeric intermediates, I_2_ and I_2_’, are most likely present at higher pHs. Similar conformational stabilities and spectroscopic properties would then appear as a single species. However, at pH below 7, the native dimer is destabilized relative to the two dimeric intermediates, and at pH 5 the first transition (N_2_⇄I_2_) disappears such that the dimeric intermediate, I_2_, is fully populated in the absence of urea. In other words, at pH 5.5 the “ native” dimer of DrPCP3b is the partially folded dimeric intermediate, I_2_, that is observed only in the presence of ∼3 M urea at higher pH. Thus, at pH 5 and pH 5.5, the equilibrium unfolding data appear as a three-state mechanism (I_2_⇄I_2_’⇄2U, see Table 1). Finally, we were not able to examine conditions below pH 5 because DrPCP3b precipitates at low pH, so we are unable to determine whether the dimer dissociates, as we have reported for HsPCP3 (6) or remains intact, as we have reported for HsPCP6 and HsPCP7 (11).

As described above (Figure 3A), the AEW of the native dimer of DrPCP3b decreased at pH<6, with an estimated pKa of ∼5.4. In addition, the equilibrium unfolding data for DrPCP3b showed that the isomerization of the native dimer (N_2_) to the dimeric intermediate (I_2_) occurs with ΔG_1_°_conf_ ∼5 kcal mol^-1^. The equilibrium unfolding data also showed that the transition is dependent on the pH (Figure 4 and Supplemental Figures S2 and S3), such that the dimeric intermediate is fully populated in the absence of urea at pH 5.5. We examined the mid-point of the transition of N_2_ to I_2_ (Figure 5A), and we fit the data as described above (Figure 3A) to estimate the pKa of the transition. The results show that the transition occurs with a pKa of ∼6.2. We suggest that the two techniques (changes in AEW and urea unfolding) likely report the same transition and that the native dimer of DrPCP3b undergoes a pH-dependent conformational change, similar to that observed in HsPCP3 and in HsPCP6, prior to unfolding. Furthermore, the data suggest that one or more active site loops are involved in the conformational change since the tryptophan residues are located in the active site.

We fit the equilibrium unfolding data globally to equation 2 (three-state; pH 5-5.5 and pH 7-8.5) or equation 3 (four-state; pH 6-6.5), and the total conformational free energies of unfolding, ΔG°_conf_, and m-values obtained from the fits over the pH range of 5 to 8.5 are summarized in Figure 5B-C and are shown in Table 1. Overall, the data show that the DrPCP3b dimer exhibits a maximum conformational free energy of ∼17 kcal mol^-1^ between pH 6 and pH 8 and that the conformational free energy decreases by ∼2 kcal mol^-1^ at higher or at lower pH. The trend is similar to that observed for HsPCP3 (6) and for HsPCP6 and HsPCP7 (11) except that HsPCP3 exhibits a larger decrease in conformational free energy (∼6 kcal mol^-1^) at higher and lower pH. The monomeric intermediate observed in HsPCP3 and in HsPCP7 is not well-populated in DrPCP3b, so it was not possible to compare the stability of the protomers for each protein. Instead, one observes that each step in unfolding of the DrPCP3b dimer contributes to the overall lower stability when compared to HsPCP3. That is, the native dimer (N_2_) as well as the dimeric intermediate (I_2_) exhibit lower stabilities compared to the same species in HsPCP3, so that collectively the total conformational free energy of DrPCP3b is ∼5 kcal mol^-1^ less than that of HsPCP3.

The cooperativity index (m-value) of each transition relates to the accessible surface area (ΔASA) exposed to solvent during unfolding (17). The results of the global fitting show that the total m-value (m_total_) is similar for DrPCP3b, HsPCP3, HsPCP6, and HsPCP7 (11). Likewise, the two observable transitions in the data for DrPCP3b exhibit similar m-values to those of HsPCP3, suggesting that similar hydrophobic surface area is exposed in the transition of N_2_ to I_2_ and in the unfolding of I_2_. In the latter case, we compared the single transition in DrPCP3b (I_2_⇄2U; ∼1 kcal mol^-1^ M^-1^) to the dimer dissociation and unfolding of HsPCP3 (I_2_⇄2I⇄2U; ∼1.7 kcal mol^-1^ M^-1^).

## Conclusions

We studied the equilibrium unfolding of procaspase-3b from *Danio rerio* in order to examine the conservation of the folding landscape of effector caspases in species other than human. We show that the constraints on the folding landscape conveyed by the requirement to maintain the caspase-hemoglobinase fold results in a conserved, yet flexible, folding landscape. The native dimer of DrPCP3b folds/unfolds similarly to those of the human effector caspases, HsPCP3, HsPCP6, and HsPCP7. The folding of DrPCP3b is more similar to that of HsPCP6 in that the monomeric intermediate (I) observed in HsPCP3 and in HsPCP7 is not well-populated. In contrast, the data for DrPCP3b show that two dimeric intermediates are well-populated at pH 6; this feature is not observed in the human effector procaspases. In addition, all effector caspases examined to date undergo a pH-dependent conformational change in the dimer, suggesting an evolutionarily conserved mechanism. Because the dimeric intermediate is enzymatically inactive (6,14), the pH-dependent transition may provide an additional mechanism for controlling enzyme activity in the cell. While the transition of N_2_ to I_2_ is reversible for all effector caspases, reforming the native dimer in the mature HsPCP3 is confounded by the irreversible dissociation of the dimer (I_2_ to 2I) at lower pH. Collectively, the data shown here for DrPCP3b and our previous data for HsPCP3, HsPCP6, and HsPCP7 (6,11) show an evolutionarily conserved folding landscape with two partially folded intermediates. Within the constraints of the conserved folding landscape, the relative population of the intermediates, as well as the overall conformational free energy, can be fine-tuned, through mutations, in a species-specific manner.

## Materials and Methods

### Reagents

IPTG and dithiothreitol (DTT) were purchased from Gold Biotechnology. Ampicillin, nickel sulfate, potassium phosphate (KH_2_PO_4_ and K_2_HPO_4_), citric acid, sodium citrate (dihydrate), tris base, imidazole, tryptone, yeast extract, and ultra-pure urea were purchased from Fisher. His-bind resin was purchased from VWR.

### Cloning, expression, and purification

Zebrafish procaspase-3a (CP-C117S) and zebrafish procaspase-3b (CP-C117S) were made by site-directed mutagenesis and were cloned into pET11a expression plasmid containing a C-terminal histidine tag, as described previously (18,19). The plasmids were transformed into *E. coli* BL21, and the proteins were purified as described (5,20).

### Phylogenetic analysis

We selected 108 sequences from the CaspBase (caspbase.org) for the phylogenetic analysis, as described previously (7). Briefly, we generated a multiple sequence alignment (MSA) in MEGA using MUSCLE (21). Both the prodomain and the intersubunit linker regions were removed. The multiple sequence alignment (MSA) was computed again using PROMALS3D (http://prodata.swmed.edu/promals3d/promals3d.php) to generate a structurally informed MSA with PDB IDs: 2j30 (human caspase-3) and 5jft (zebrafish caspase-3a) (22). The phylogenetic tree was generated using IQTREE using the Jones-Taylor Thornton model (JTT) plus gamma distribution (23), and the tree was bootstrapped 1000 times as a test of phylogeny.

### Stock solutions

Urea stock solutions (10 M) were made as described previously (15) in citrate buffer (50 mM sodium citrate/citric acid, pH 5-6, 1 mM DTT), phosphate buffer (50 mM potassium phosphate monobasic/potassium phosphate dibasic, pH 6.5-8, 1 mM DTT), or Tris-HCl buffer (50 mM Tris-HCl, pH 8.5, 1 mM DTT). All solutions were prepared fresh for each experiment and were filtered (0.22 μm pore size) prior to use. The urea stock solutions were prepared by weight and the concentration was examined by refractive index (15), and solutions were used if the two values were within ±1%.

### Size exclusion chromatography

In separate experiments, DrPCP3b (CP-C117S) was dialyzed for ∼16 hours at 25 °C in a buffer of 50 mM phosphate (pH 6 to pH 8), 50 mM Tris-HCl (pH 8.5), or 50 mM citrate buffer (pH 5 to pH 6). The protein was diluted to a concentration of 25 μM. Protein (100 μL) was loaded onto a Superdex75 Increase 10/300GL column that had been pre-equilibrated with the dialysis buffer. The protein was eluted using a flow rate of 0.8 mL min^-1^ on an AKTA FPLC system with UPC-900 Detector and P920 pump. The absorbance of the eluant was measured at 280 nm, and the column was standardized using the gel filtration LMW calibration kit (GE Health Sciences, 28-4038-41), following the manufacturer’s instructions.

### Equilibrium unfolding

DrPCP3b was dialyzed in citrate buffer for experiments from pH 5 to pH 6, phosphate buffer for experiments from pH 6.5 to pH 8, and Tris-HCl buffer for experiments at pH 8.5. All buffers contained 1 mM DTT. Equilibrium unfolding experiments were performed as described (5). Briefly, in each pH, the protein concentrations were varied from 0.5 to 8 μM. To confirm that folding is reversible, the protein was first incubated in 9 M urea-containing buffer for 4 hours, then the samples were diluted to the urea concentrations shown in the figures and equilibrated for at least 16 hours. All samples were incubated at 25 °C for a minimum of 16 hours. For each sample, fluorescence emission was acquired from 300-400 nm following excitation at 280 or 295 nm (PTI C-61 spectrofluorometer, Photon Technology International). Circular dichroism was measured at 232 nm for pH 5 and pH 5.5 and 224 nm for pH between pH 6 and pH 8.5 with a Jasco J1500 spectropolarimeter.

### Data analysis

For equilibrium folding/unfolding studies at multiple pH values, the average emission wavelength for each fluorescence emission scan was calculated using equation 1,

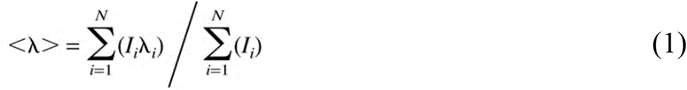

where <λ> is the average emission wavelength (AEW), and *I*_i_ is the fluorescence emission at wavelength λ_i_ (24).

Data collected for pH 5 and pH 5.5, as well as between pH 7 to pH 8.5, were fit to a 3-state equilibrium folding model as described previously (6) and shown in equation 2.

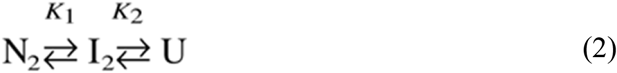

In this model, the native dimer (N_2_) isomerizes to a partially folded intermediate dimer (I_2_), which then unfolds to the monomer (2U). The equilibrium constants, *K*_1_ and *K*_2_, correlate to the two unfolding steps, respectively.

Data collected at pH 6 and pH 6.5 were fit to a four-state equilibrium folding model as described (6) and shown in equation 3.

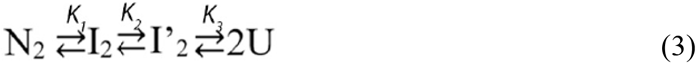

In this model, the native dimer (N_2_) isomerizes to a dimeric intermediate (I_2_), and the dimeric intermediate isomerizes to a second partially folded dimer (I_2_’) prior to unfolding (2U). The equilibrium constants, *K*_1_, *K*_2,_ and *K*_3_, correlate to the three unfolding steps, respectively.

### Global fits of the equilibrium unfolding data

The experimental protocol results in 10-12 data sets at each pH, corresponding to three spectroscopic probes and three to four protein concentrations, as described above. The data collected at each pH were fit (Igor Pro) using global fitting to the three-state (equation 2) or four-state (equation 3) models described above and as described previously (15). Briefly, the global fits yield the free energy changes in the absence of denaturant, corresponding to the equilibrium constants *K*_1_, *K*_2_ and *K*_3_, respectively, as well as the cooperativity indices, *m*_1_, *m*_2_, *m*_3_, associated with each step of unfolding, as described previously (15,20). The results of the global fits at each pH are shown as the solid lines in the figures, and the values are presented in Table 1.

## ACKNOWLEDGEMENTS

This work was supported by a grant from the National Institutes of Health [grant number GM127654 (to A.C.C.)] and by funds from UT Arlington [Office of the Vice President for Research (to A.C.C.)].

