## Supplemental Figures Tables for "Comparing the folding landscapes of evolutionarily divergent procaspase-3"

**Running title**: Evolutionarily conserved caspase folding landscape

[*]Corresponding author: A. Clay Clark

**Key Words**: caspase; dimerization; apoptosis; protein folding; florescence spectroscopy; circular dichroism; zebrafish

**Supplemental Table S1. Genes selected for phylogenetic tree.**

| Accession ID | Speicies name | Caspase type |
|---|---|---|
| NP_001217.2 | Homo sapiens | 6 |
| NP_001253985.1 | Homo sapiens | 7 |
| NP_004337.2 | Homo sapiens | 3 |
| NP_001012435.1 | Pan troglodytes | 3 |
| XP_016774869.1 | Pan troglodytes | 7 |
| XP_016807502.1 | Pan troglodytes | 6 |
| XP_002815098.1 | Pongo abelii | 6 |
| XP_003777685.1 | Pongo abelii | 7 |
| XP_009238771.1 | Pongo abelii | 3 |
| NP_037054.1 | Rattus norvegicus | 3 |
| NP_071596.1 | Rattus norvegicus | 7 |
| NP_113963.2 | Rattus norvegicus | 6 |
| NP_001018333.1 | Danio rerio | 6 |
| NP_001018443.1 | Danio rerio | 7 |
| NP_571952.1 | Danio rerio | 3 |
| XP_005173133.1 | Danio rerio | 3 |
| XP_005156389.1 | Danio rerio | 7 |
| XP_001513388.4 | Ornithorhynchus anatinus | 7 |
| XP_001517122.2 | Ornithorhynchus anatinus | 3 |
| NP_001157433.1 | Equus caballus | 3 |
| XP_005607953.1 | Equus caballus | 6 |
| XP_014588814.1 | Equus caballus | 7 |
| NP_990056.1 | Gallus gallus | 3 |
| NP_990057.1 | Gallus gallus | 6 |
| XP_421764.3 | Gallus gallus | 7 |
| NP_001011068.1 | Xenopus tropicalis | 6 |
| NP_001016299.1 | Xenopus tropicalis | 7 |
| XP_007905080.1 | Callorhinchus milii | 3 |
| XP_007907939.1 | Callorhinchus milii | 6 |
| XP_007422761.1 | Python bivittatus | 3 |
| XP_007437451.1 | Python bivittatus | 6 |
| XP_007441954.1 | Python bivittatus | 7 |
| XP_005999642.1 | Latimeria chalumnae | 6 |
| XP_006002865.1 | Latimeria chalumnae | 7 |
| XP_014351567.1 | Latimeria chalumnae | 3 |
| XP_004549328.1 | Maylandia zebra | 3 |

| | | |
|---|---|---|
| XP_004563667.1 | Maylandia zebra | 3 |
| XP_004567546.1 | Maylandia zebra | 7 |
| XP_014268841.1 | Maylandia zebra | 6 |
| XP_005797675.2 | Xiphophorus maculatus | 6 |
| XP_005804467.1 | Xiphophorus maculatus | 7 |
| XP_005807677.1 | Xiphophorus maculatus | 3 |
| XP_005282031.1 | Chrysemys picta bellii | 3 |
| XP_005287972.1 | Chrysemys picta bellii | 6 |
| XP_005293940.1 | Chrysemys picta bellii | 7 |
| XP_005044995.1 | Ficedula albicollis | 6 |
| XP_005045177.1 | Ficedula albicollis | 3 |
| XP_005048827.1 | Ficedula albicollis | 7 |
| XP_003823510.1 | Pan paniscus | 3 |
| XP_003825647.1 | Pan paniscus | 7 |
| XP_003830063.1 | Pan paniscus | 6 |
| XP_004639834.1 | Octodon degus | 3 |
| XP_005406246.1 | Chinchilla lanigera | 6 |
| XP_013368516.1 | Chinchilla lanigera | 3 |
| XP_013371286.1 | Chinchilla lanigera | 7 |
| XP_004426643.1 | Ceratotherium simum simum | 6 |
| XP_004428794.1 | Ceratotherium simum simum | 3 |
| XP_014641239.1 | Ceratotherium simum simum | 7 |
| XP_006831548.1 | Chrysochloris asiatica | 7 |
| XP_006834493.1 | Chrysochloris asiatica | 3 |
| XP_006869403.1 | Chrysochloris asiatica | 6 |
| NP_001098140.1 | Oryzias latipes | 3 |
| NP_001098168.1 | Oryzias latipes | 3 |
| XP_004080311.2 | Oryzias latipes | 7 |
| XP_004084686.1 | Oryzias latipes | 6 |
| XP_005500235.1 | Columba livia | 3 |
| XP_005515276.1 | Columba livia | 6 |
| XP_021151155.1 | Columba livia | 7 |
| XP_006022892.1 | Alligator sinensis | 7 |
| XP_006025610.2 | Alligator sinensis | 6 |
| XP_006026683.1 | Alligator sinensis | 3 |
| XP_009879870.1 | Charadrius vociferus | 7 |
| XP_009883787.1 | Charadrius vociferus | 6 |
| XP_009890080.1 | Charadrius vociferus | 3 |
| XP_010000752.1 | Chaetura pelagica | 3 |

| | | |
|---|---|---|
| XP_010002388.1 | Chaetura pelagica | 6 |
| XP_010006222.1 | Chaetura pelagica | 7 |
| XP_017263180.1 | Kryptolebias marmoratus | 6 |
| XP_017278585.1 | Kryptolebias marmoratus | 3 |
| XP_017289362.1 | Kryptolebias marmoratus | 7 |
| NP_001081225.1 | Xenopus laevis | 3 |
| NP_001081406.1 | Xenopus laevis | 6 |
| NP_001081408.1 | Xenopus laevis | 7 |
| NP_001091272.1 | Xenopus laevis | 7 |
| XP_019389662.1 | Crocodylus porosus | 6 |
| XP_019390229.1 | Crocodylus porosus | 3 |
| XP_019411759.1 | Crocodylus porosus | 7 |
| XP_019358381.1 | Gavialis gangeticus | 7 |
| XP_019377452.1 | Gavialis gangeticus | 3 |
| XP_019377505.1 | Gavialis gangeticus | 6 |
| NP_001269823.1 | Oreochromis niloticus | 3 |
| XP_003451815.1 | Oreochromis niloticus | 3 |
| XP_003451816.1 | Oreochromis niloticus | 3 |
| XP_003453649.2 | Oreochromis niloticus | 7 |
| XP_003453650.2 | Oreochromis niloticus | 3 |
| XP_013124898.1 | Oreochromis niloticus | 7 |
| XP_019215518.1 | Oreochromis niloticus | 6 |
| XP_019215520.1 | Oreochromis niloticus | 6 |
| XP_020447497.1 | Monopterus albus | 6 |
| XP_020451082.1 | Monopterus albus | 3 |
| XP_020460887.1 | Monopterus albus | 7 |
| XP_021381152.1 | Lonchura striata domestica | 6 |
| XP_021381698.1 | Lonchura striata domestica | 7 |
| XP_021399050.1 | Lonchura striata domestica | 3 |

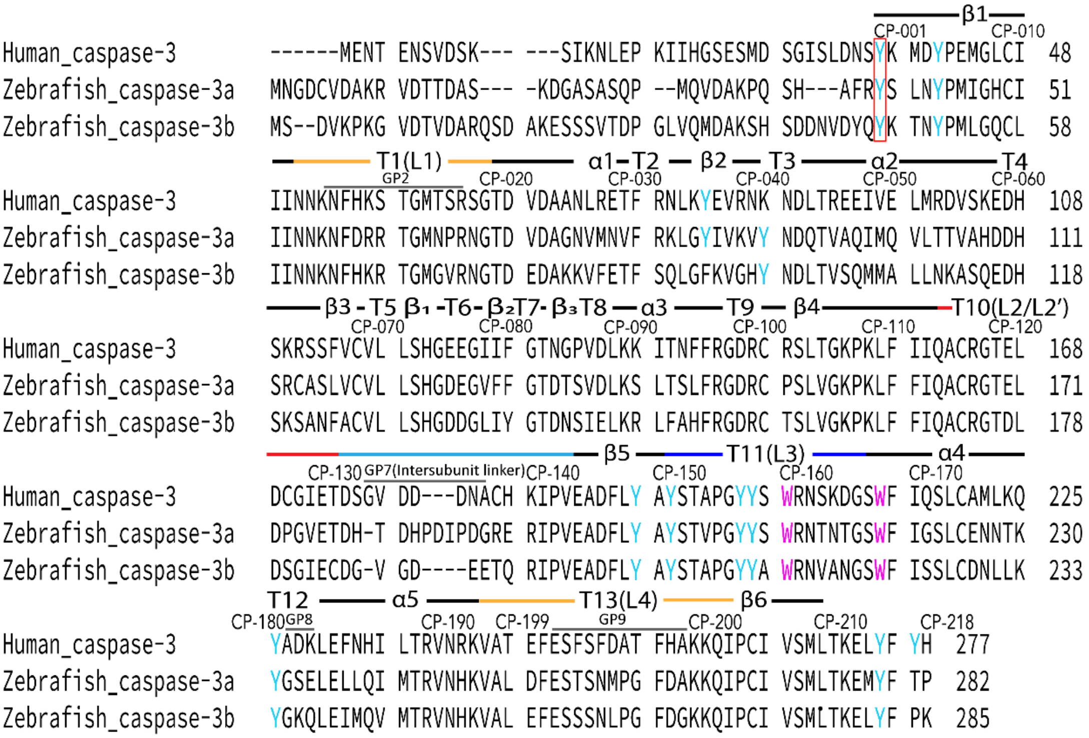


**Supplemental Figure S1. Comparison of sequences for HsPCP3, DrPCP3a and DrPCP3b.** The common position numbering scheme, as described in the text, and secondary structural elements are shown above each line, and actual position numbers are shown at the end of each row. The tryptophan and tyrosine residues are marked as purple and cyan, respectively.

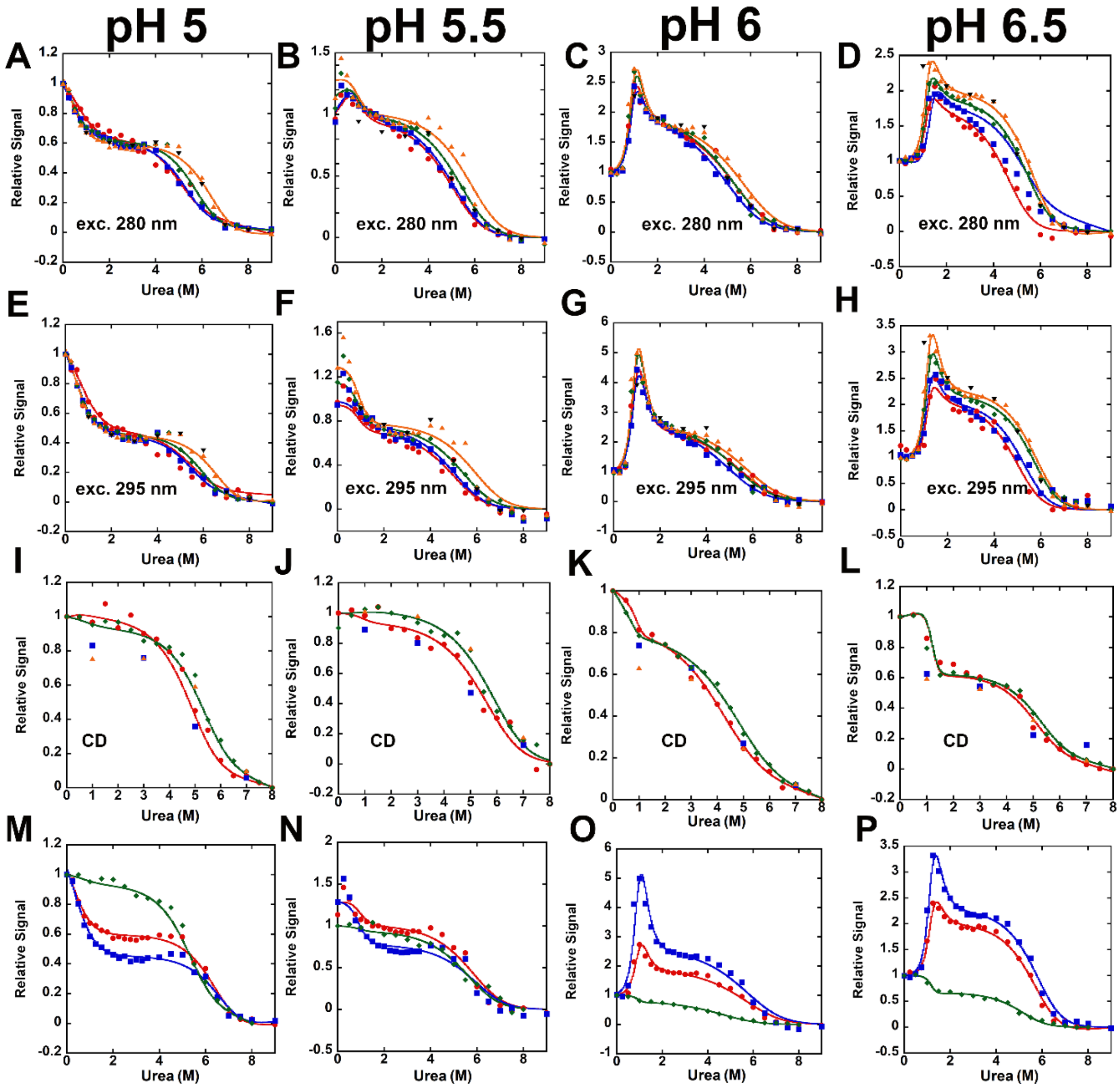


**Supplemental Figure S2. Equilibrium unfolding versus urea of zebrafish procaspase-3b (CP-C117S) (called DrPCP3b) from pH 5 to pH 6.5.** The pH is indicated on the top of each column. For panels A-H, the protein concentrations are as follows: 0.5 µM (●), 1 µM (■), 2 µM (♦) and 4 µM (▲). For panels I-L, the protein concentrations are as follows: 2 µM (●), 4 µM (♦). Refolding data demonstrates reversible folding/unfolding at each pH; for panels A-H, refolding data for 4 µM protein concentration are shown (▼). For panels I-L, refolding data for 2 µM (■), 4 µM (▲) are shown. For panels A-H, unfolding was monitored by fluorescence emission with excitation at 280 nm (panels A-D) or 295 nm (panels E-H). For panels I-J, unfolding was monitored by circular dichroism at 224 nm or 232 nm. Panels M-P show a comparison of unfolding for 4 µM protein concentration and fluorescence emission with excitation at 280 nm (●) or 295 nm (■) and CD (♦). The solid lines in each panel represent global fits of the data as described in the text.

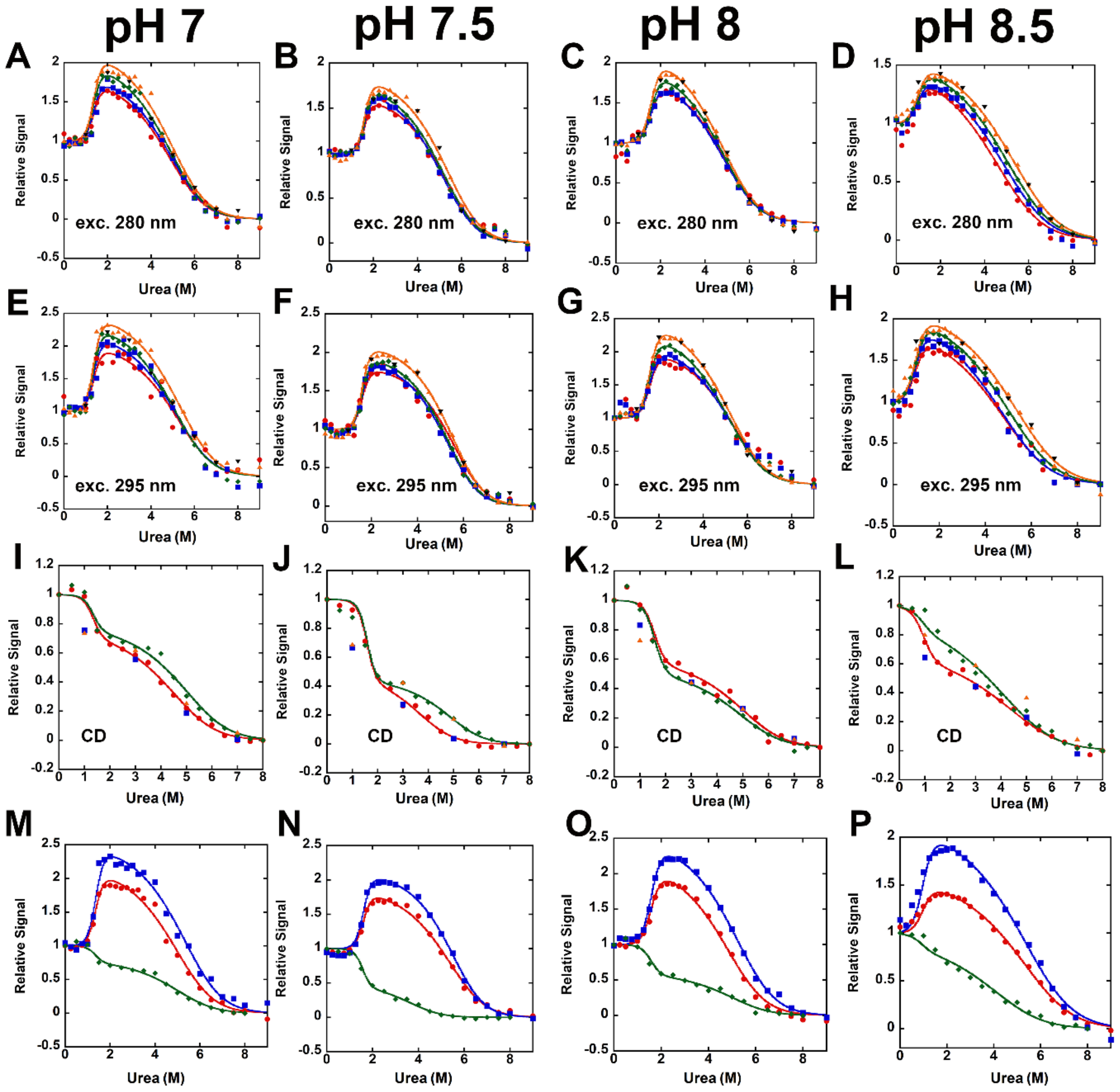


**Supplemental Figure S3. Equilibrium unfolding versus urea of zebrafish procaspase-3b (CP-C117S) (called DrPCP3b) from pH 7 to pH 8.5.** The pH is indicated on the top of each column. For panels A-H, the protein concentrations are as follows: 0.5 µM (●), 1 µM (■), 2 µM (♦) and 4 µM (▲). For panels I-J, the protein concentrations are as follows: 2 µM (●), 4 µM (♦). For panels K-L, the protein concentrations are as follows: 4 µM (●), 8 µM (♦). Refolding data demonstrates reversible folding/unfolding at each pH; for panels A-H, refolding data for 4 µM protein concentration are shown (▼). For panels I-L, refolding data for 2 µM (■), 4 µM (▲) are shown. For panels A-H, unfolding was monitored by fluorescence emission with excitation at 280 nm (panels A-D) or 295 nm (panels E-H). For panels I-J, unfolding was monitored by circular dichroism at 224 nm or 232 nm. Panels M-P show a comparison of unfolding for 4 µM protein concentration and fluorescence emission with excitation at 280 nm (●) or 295 nm (■) and CD (♦). The solid lines in each panel represent global fits of the data as described in the text.
